# FM-dye inhibition of Piezo2 relieves acute inflammatory and osteoarthritis knee pain in mice of both sexes

**DOI:** 10.1101/2025.03.17.643683

**Authors:** Natalie S. Adamczyk, Shingo Ishihara, Alia M. Obeidat, Dongjun Ren, Richard J. Miller, Anne-Marie Malfait, Rachel E. Miller

**Affiliations:** Rush University Medical Center, Department of Internal Medicine, Division of Rheumatology, Chicago, IL, USA; Chicago Center on Musculoskeletal Pain, Chicago, IL, USA; Northwestern University, Department of Pharmacology, Chicago, IL, USA

**Keywords:** Osteoarthritis, Piezo2, pain, nociceptor, sensitization

## Abstract

Musculoskeletal pain is a significant burden affecting billions of people with little progress in the development of pharmaceutical pain relief options. The mechanically-activated ion channel Piezo2 has been shown to play a role in mechanical sensitization; however there has been little progress in examining therapeutics that target this molecule. The goal of this study was to assess the effect of two FM-dyes, FM1-43 or FM4-64, in reducing acute inflammatory and osteoarthritis knee joint pain in mice of both sexes. In our acute model of Complete Freund’s adjuvant (CFA)-induced joint pain, mice intra-articularly injected with FM1-43 exhibited an attenuation of knee hyperalgesia 90 minutes following injection. *In vivo* calcium imaging of the dorsal root ganglion (DRG) also demonstrated a reduction in nociceptor responses to mechanical forces applied to the knee joint of CFA mice following FM-dye injection. Male and female WT mice subjected to partial medial meniscectomy (PMX) surgery as a model of osteoarthritis developed more severe knee hyperalgesia than nociceptor-specific Piezo2 conditional knock-out mice. Intra-articular injection of FM1-43 reduced both knee hyperalgesia and weight-bearing asymmetry in this model and had no effect in Piezo2 conditional knock-out mice. Finally, in mice with spontaneous osteoarthritis associated with aging, intra-articular injection of FM-dyes also reduced knee hyperalgesia. In conclusion, inhibiting Piezo2 genetically or pharmacologically was effective in reducing pain-related behaviors in mice of both sexes in the setting of inflammatory and osteoarthritis knee pain. These studies provide evidence of the therapeutic potential of targeting Piezo2 in musculoskeletal pain conditions.

## Introduction

Globally 1.71 billion people suffer from musculoskeletal ailments ranging from rheumatologic to orthopedic pain conditions [6]. Osteoarthritis (OA) is among the top 10 leading causes of disability worldwide, affecting women more often than men [28]. Individuals with OA often suffer from pain associated with movement and mechanical loading, such as stair climbing and sit-to-stand tasks, indicating knee OA is highly mechanosensitive in nature [10,11,28].

The key cell type responsible for sensing noxious stimuli in the peripheral nervous system is the nociceptor. Expression of ion channels enables nociceptors to respond to various chemical, thermal and mechanical stimuli. In pathological states such as OA, nociceptors become sensitized, resulting in amplified pain signals and the interpretation of normally innocuous stimuli as painful. This peripheral sensitization is a hallmark of OA pain and contributes to the development of chronic pain. Piezo2 is a nonselective cation channel that has been shown to play a role in the mechanical sensitization of nociceptors when exposed to inflammatory conditions [9,27,33]. We have previously demonstrated that genetic depletion of Piezo2 in nociceptors resulted in reduced mechanical sensitization in models of inflammatory and OA knee pain [31]. However, whether targeting Piezo2 pharmacologically could prove beneficial in reducing mechanically-evoked pain in conditions like OA has yet to be tested.

To date there are no specific Piezo2 agonists or antagonists, and published work has focused on increasing cell membrane stiffness or utilizing nonspecific inhibitors such as margaric acid or GsMTx-4 [2,33,35]. Importantly, a recent report demonstrated that the cationic styryl dye, FM1-43, and its derivative FM4-64, are selective for Piezo2 *in vivo* by demonstrating that the dye is not taken up by nerve terminals in Piezo2 conditional knock-out mice [36]. It has been shown that intraplantar injection of FM1-43 can increase mechanical withdrawal thresholds in mice [8,12], and evidence suggests that inhibitory actions of FM1-43 are produced as the dye passes through the pore competing with cations for binding sites [13,24].

Therefore, the goal of the current study was to assess the inhibitory effect of FM-dyes in acute inflammatory and chronic osteoarthritis knee joint pain in mice of both sexes, and to determine whether this effect is mediated through Piezo2. We also aimed to determine whether FM-dye injected into the knee joint is acting on nociceptors by assessing its effects during *in vivo* calcium imaging of the DRG.

## Methods

### Animals

Mice were housed in a climate-controlled animal care facility at Rush University Medical Center receiving food (Teklad Global 18% Protein Rodent Diet, Inotiv #2018) and water *ad libitum*. Mice were on a 12hr light cycle from 7am-7pm. All procedures were approved by the Rush and Northwestern University IACUC committees. Na_V_1.8-cre^+/-^;tdTomato^fl/+^;Piezo2^fl/fl^ (Piezo2cko), and Na_V_1.8-cre^+/-^;GCaMP6s^fl/+^ (Na_V_1.8-GCaMP6) mice were bred at Rush University Medical Center. Age-matched wild type (WT) C57BL/6 control mice were obtained from Jackson Laboratories. For aging experiments, Na_V_1.8-cre;GCaMP6s-loxp;Piezo2-loxp or Na_V_1.8-cre;tdTomato-loxp;Piezo2-loxp mice of mixed genotypes were bred and aged in house as described below. For all experiments, cages were randomized to experimental condition.

### FM-dye injections

FM1-43FX (Invitrogen F35355) and FM4-64FX (Invitrogen F34653) were prepared by dissolving the dyes in sterile saline. FM1-43 and FM 4-64 are closely related structurally, with slight differences that result in different fluorescence emission spectra: FM1-43 emits in the GFP range, while FM4-64 emits in the RFP range [3]. The uptake of both dyes has previously been shown to be dependent on Piezo2 activity[36]. Therefore, we chose to use FM1-43 in mice that would be expressing tdTomato, and FM4-64 in mice expressing GCaMP6 in order to avoid interference based on the FM-dye emission.

Based on prior literature, which showed efficacy two hours following an intra-plantar injection of 5 nmol FM1-43 to the hind paw [8,12], we first performed a dose-response study to assess the effective dose for intra-articular (i.a.) injections and the proper testing time point. We injected 0, 0.05, 0.5, 5, or 50 nmol of FM1-43 in 2.5 µL into the right knees of WT male mice three days post Complete Freund’s adjuvant (CFA) injection, and measured knee hyperalgesia before, and 0.5, 1.5, 3, 6, and 24 hours after injection (n=2 mice/dose). Both the 5 and 50 nmol conditions reduced knee hyperalgesia, with a maximum reversal observed 90 min post FM injection (Supplementary Figure 1), which was consistent with the prior literature [8,12]. From this experiment, we decided to use an i.a. dose of 5 nmol in 2.5 µL and to evaluate mice 90 minutes post FM injection for all subsequent behavior experiments.

### Complete Freund’s adjuvant (CFA) model of acute inflammatory knee pain

CFA (Millipore, 344289) (5 µL stock, n=4 female; n=4 male) or saline vehicle (5 µL, n=4 female; n=4 male) was intra-articularly (i.a.) injected into the right knees of 12-week old WT mice of both sexes, as previously described [31]. Briefly, mice were placed under isoflurane anesthesia and the right knee was shaved and the patellar tendon was visually located. A sub-patellar injection was performed with a 30 G needle without any incision. Knee width was measured using a caliper. To study knee hyperalgesia, FM-dye or vehicle injections began 3 days after model induction.

### PMX Surgery Model of OA Knee Pain

Partial medial meniscectomy (PMX) surgery was performed in male and female 12-week old WT (n=4/sex for knee hyperalgesia; n=4 female and n=5 male for weight-bearing asymmetry) or Piezo2cko (n=4/sex for knee hyperalgesia) mice as previously described [19,29]. WT sham-operated mice were also used as a control (n=4/sex for knee hyperalgesia; n=4 female and n=5 for weight-bearing asymmetry). Mice were placed under isoflurane anesthesia (1-1.5% in O_2_) and their right knees were shaved and sanitized with betadine. The joint capsule was opened, the medial meniscotibial ligament was severed with a scalpel, and approximately 1/3 of the medial meniscus was removed to induce joint destabilization. Sham surgery consisted of the joint capsule being opened. In both cases, the joint capsule was closed via suture followed by closure of the skin incision with suturing. No analgesics were provided post-operatively.

To study knee hyperalgesia, n=4 WT and Piezo2cko mice of each sex were used for PMX or sham surgery, and FM-dye or vehicle injections began 8 weeks after surgery. On the first injection day, mice were assessed for knee hyperalgesia (pre-injection reference). Following this test, mice were randomly assigned to receive either vehicle or FM1-43 i.a. injection and assessed again for knee hyperalgesia 90 minutes later. In order to reduce the number of animals needed for this experiment in part due to the limited availability of Piezo2cko mice, we utilized a cross-over design. Therefore, 4 days later mice were re-tested to establish a pre-injection withdrawal threshold, and mice switched conditions such that those that originally received FM now received vehicle and vice versa. Since the pre-injection values were similar for individual mice regardless of the order of injection, data was combined for statistical analysis (Supplemental Figure 2A-C).

To study weight bearing asymmetry, n=4 WT male mice and n=5 WT female mice were used for PMX or sham surgery, and FM-dye or vehicle i.a. injections began 12 weeks after surgery. Again, to reduce the numbers of mice needed, 2 days later mice were re-tested to establish a pre-injection baseline, and mice switched conditions such that those that originally received FM now received vehicle and vice versa. Since the pre-injection values were similar for individual mice regardless of the order of injection, data was combined for statistical analysis. Following weight bearing asymmetry testing, these mice were also used for horizontal ladder testing 2 days later.

### Aging model

For studying the effects of Piezo2 depletion in nociceptors with age, Na_V_1.8-cre;GCaMP6s-loxp;Piezo2-loxp or Na_V_1.8-cre;tdTomato-loxp;Piezo2-loxp mice were bred such that the offspring would contain a mixture of genotypes. Mice that either did not have any copies of Na_V_1.8-Cre present or did not have any copies of floxed Piezo2 present were considered ‘WT’. Mice that had one or two copies of Na_V_1.8-cre and one or two copies of Piezo2-loxp were considered ‘Piezo2cko’ for this experiment, meaning that both Piezo2cko heterozygous and Piezo2cko homozygous mice were used, as we have previously shown that both heterozygous and homozygous Piezo2cko mice had a reduction in Piezo2 levels in Na_V_1.8+ neurons [31]. In total, naïve WT and Piezo2cko littermates of both sexes (Male: WT n=4, Piezo2cko n=6; Female: WT n=6, Piezo2cko n=7) were left to age for 24 months, at which time knee hyperalgesia was evaluated following i.a. injections of either FM1-43 or FM4-64 depending on if the genotype was tdTomato or GCaMP6s, respectively.

### Knee Hyperalgesia

Knee hyperalgesia was assessed as previously described [25,31]. Mice were manually restrained with the knee in flexion, and an experimenter blinded to condition was guided by software to increase pressure on the right knee using an Ugo Basile PAM device. Once a pain response was elicited (vocalization, head jerk, knee jerk, or muscle spasm), the trial was stopped, and the responding force was recorded. If no pain response was elicited, the mouse received a maximum score of 450 g. Two trials were conducted and then averaged to calculate the knee withdrawal threshold.

### Weight bearing asymmetry

Weight bearing asymmetry was assessed as previously described [31]. Briefly, one week prior to testing mice were habituated to the task in a cage void of stimulus other than 20 strings of various lengths half of which were baited with a Honey Nut Cheerio. After one hour, any mouse pulling fewer than 15/20 strings were retrained within the same week. If the mice still failed after retraining, they were dropped from the study; in this study one such mouse was dropped. The week following training, mice were tested by a blinded individual for weight bearing asymmetry utilizing a Bioseb static incapacitance device with a custom-built plexiglass open box chamber (65mm x 140mm x 260mm). During the testing, a Cheerio baited string was draped over the top of the chamber to assist in positioning the mouse to stand with one leg on each force sensor. When the mouse was positioned appropriately and pulling on the string, a weight-bearing measurement was taken with forces averaged over 2 seconds. Mice were measured three times, and an asymmetry score (right leg grams - left leg grams) was calculated and averaged from the three trials.

### Horizontal Ladder

Mice were placed on a horizontal ladder (60 cm in length) with evenly spaced bars (2 cm apart) [1,23]. Mice were run across the ladder once and the number of slips of the right hind leg was calculated from the beginning to the end point of the task from a video recording by a blinded individual.

### In vivo DRG calcium imaging

*In vivo* calcium imaging of the L4 DRG was conducted as previously described [18,26,31]. Na_V_1.8-GCaMP6 mice (n= 8 female; n= 5 male) were i.a. injected with CFA. One day prior to imaging mice were shipped to Northwestern University for *in vivo* two-photon imaging of the L4 DRG. All procedures were approved by Northwestern’s IACUC. Three days post injection of CFA, mice were anesthetized with isoflurane anesthesia (1.5–2% in O_2_) and underwent a dorsal laminectomy to expose the right-side L4 DRG. For the remainder of the imaging session, mice were maintained under isoflurane anesthesia. To prevent the L4 DRG from drying, Silicone elastomer (World Precision Instruments) was applied to the exposed tissue. A custom stage with Narishige spinal clamps was used to elevate the mouse to prevent breathing artifacts. Once the mouse was elevated, the stage was positioned under a Prairie Systems Ultima In Vivo two photon microscope. Image acquisition was controlled using PrairieView software version 5.3 or version 5.5 with a Coherent Chameleon-Ultra2 Ti:Sapphire laser tuned to 920 nm. Prior to any stimulus being applied the mouse was imaged for 30 frames as a baseline. A calibrated forceps instrument was used to apply a mechanical stimulus of 100g to the medial-lateral aspect of the mouse knee joint during imaging. After the first mechanical stimulus was applied, mice were injected with FM4-64 or vehicle (6 nmol in 3 µL) into the right knees. After 90 minutes had passed, 100g of force was applied again to the knee. For each 100g stimulus recording, the first 10 frames served as a baseline, during the second set of 10 frames the mechanical stimulus was applied, and the final 10 frames served as a post stimulus baseline. To analyze the videos a custom ImageJ macro was used to quantify the changes in fluorescence over time (https://mskpain.center/download_file/8dd82cc8-6c1c-4e37-b466-279afabd6307/530) using the formula ΔF/Fo, where Fo is the average intensity of the baseline period acquired without force. The number of neurons responding to the force before and after the FM or vehicle injection were manually identified by an individual blinded to conditions. Based on the area of the DRG imaged and the prior calculated density of neurons, the total number of neurons imaged was extrapolated [26,31]. Data were represented as the number of responding neurons / total number of neurons imaged *100. Representative heatmaps depicting the ΔF/Fo for each responding neuron (column) for each video frame (row) were generated using GraphPad Prism.

### Knee Histology

Right knees from the weight-bearing experiment were placed in Zamboni’s fixative overnight and washed the next day in DI water. Knees were decalcified for two weeks in 14% EDTA and placed in sucrose until the tissue sunk. Knees were frozen in OCT on dry ice and sectioned using a Cryostat with a CryoJane system at 20 µm. Mid-joint knee sections were used for Safranin-O or H&E staining, as previously described [29]. Safranin-O was used to quantify cartilage degeneration, and H&E stained sections were used to score osteophyte width and synovitis by a person blinded to treatment, as previously described [29,30].

### Statistics

All statistical analysis was performed in GraphPad Prism version 9 or 10. For behavioral data, male and female results were pooled for analyses – individually labeled data points are graphed for comparison purposes. For histological data, male and female results were evaluated separately due to the known difference in structural joint damage that develops in male and female mice [21,22]. For the CFA model, a two-way repeated measures ANOVA with Sidak’s multiple comparisons test was used to assess knee hyperalgesia and knee swelling between saline and CFA mice at baseline and at day 3 prior to FM injections; to compare a single condition’s (CFA or saline treated) knee withdrawal response pre and post either vehicle or FM1-43, a paired two-tailed t-test was utilized. For *in vivo* calcium imaging, a paired two-tailed t-test was used to compare the percentage of responding neurons before and after vehicle or FM4-64 injection. For PMX knee hyperalgesia, an ordinary one-way ANOVA with Tukey’s post hoc test was used to analyze the difference between WT sham, WT PMX and Piezo2cko PMX 8 weeks after surgery for differences in knee hyperalgesia. As with the CFA study, to compare a single group’s response pre and post either FM1-43 or vehicle, a paired two-tailed t-test was utilized. For PMX weight-bearing, an unpaired two-tailed Student’s t-test was used to assess differences between WT sham and PMX mice for weight-bearing asymmetry, paired two-tailed t-tests were used to assess a single group’s response pre and post either FM1-43 or saline. An unpaired two-tailed Student’s t-test was used to compare aged mouse knee hyperalgesia between WT and Piezo2cko mice, and a paired two-tailed t-test was used to assess pre vs post response to FM. Unless otherwise indicated, mean and standard error of the mean (SEM) are presented.

## Results

### Injection of FM1-43 alleviates inflammatory knee pain in wild-type mice of both sexes

To assess the effectiveness of FM1-43 in alleviating acute inflammatory knee pain, we used the previously validated CFA intra-articular injection model [4,31]. Baseline knee withdrawal and knee width measurements were established prior to induction of the model, and no differences were observed in mice allocated to the CFA or saline groups (Figure 1A-C). By the morning of day 3, male and female mice injected with CFA had a reduced knee withdrawal threshold compared to mice injected with saline (p<0.0001, Figure 1B). Mice injected with CFA also had increased knee width by day 3 compared to saline controls (p<0.0001, Figure 1C). CFA and saline-injected mice were then allocated to vehicle or FM1-43 treatment groups. Treatments were intra-articularly injected in the afternoon of day 3, and mice were evaluated for knee withdrawal threshold 90 minutes after the injection. CFA mice injected with FM1-43 had an increased knee withdrawal threshold compared to their morning test, indicating a reduction in knee hyperalgesia (pre 287±12 g; post 406±3 g; p=0.0017, Figure 1D). In contrast, CFA mice injected with vehicle had a trend toward a reduced withdrawal threshold from their morning test, indicating that they developed more knee hyperalgesia (pre 331±15 g; post 305±8 g; p=0.0759, Figure 1E). While saline mice had not developed knee hyperalgesia by day 3 (Figure 1B), injection with FM1-43 slightly increased the knee withdrawal threshold (pre 414±3 g; post 428±3 g; p=0.0026, Figure 1F), while vehicle injection slightly lowered the knee withdrawal threshold (pre 419±2 g; post 407±4 g; p= 0.0991, Figure 1G). Together these data suggest that FM1-43 injected locally can reduce acute inflammatory knee pain with minimal effects observed when injected in a healthy knee joint.

**Figure 1.**
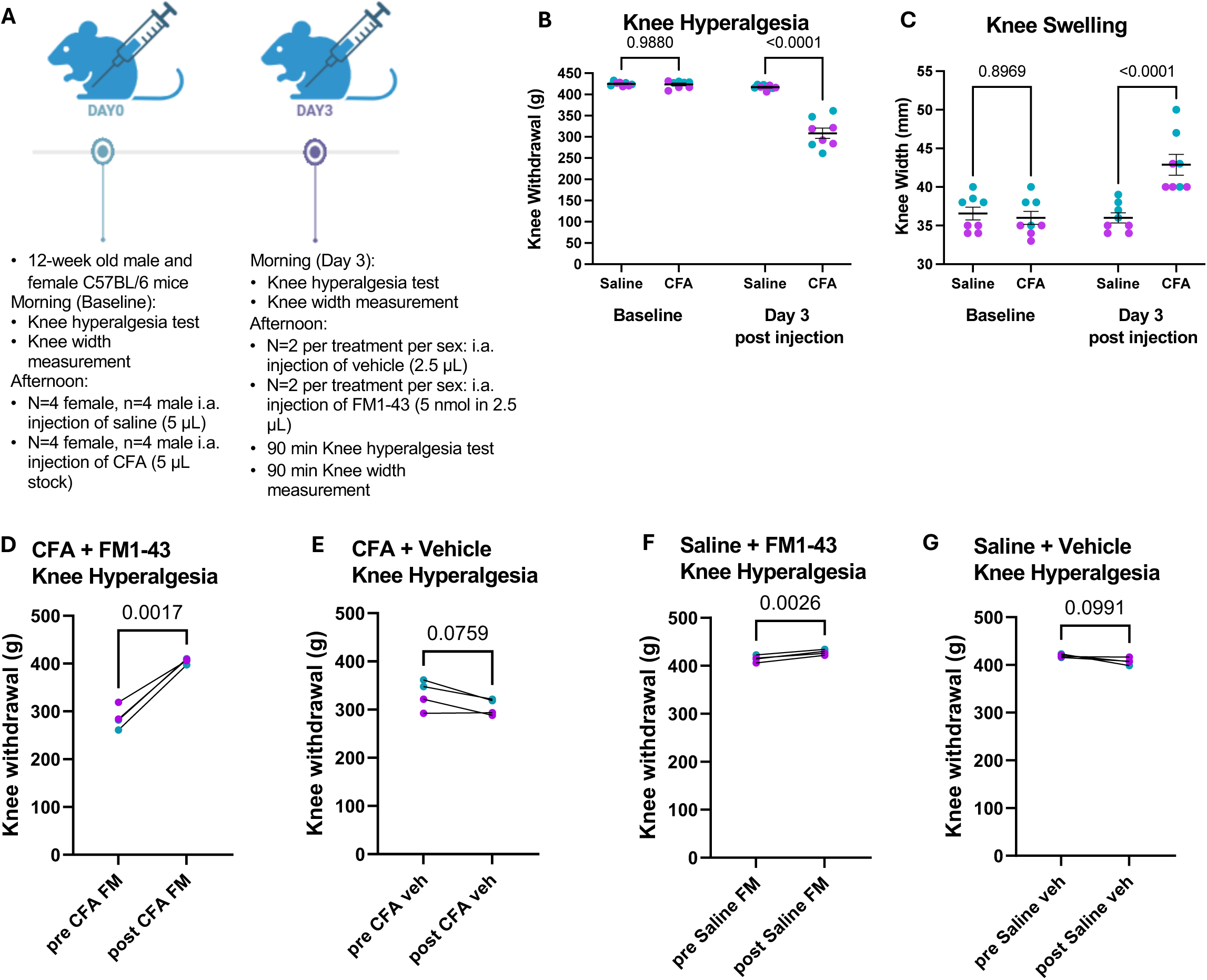
Intra-articular injection of FM1-43 reverses knee hyperalgesia in the CFA model of acute knee pain. A) Experimental timeline. B) Knee withdrawal threshold before and 3 days after i.a. injection of CFA (5 μL of stock) or vehicle. Two-way repeated measures ANOVA with Sidak’s multiple comparisons test. C) Knee width measurement of the mice before and 3 days after i.a. injection of CFA or vehicle. Two-way repeated measures ANOVA with Sidak’s multiple comparisons test. D-G) Mice in the CFA and Vehicle conditions were sub-divided into vehicle and FM1-43 groups. Knee withdrawal threshold numbers before the second injection are marked here as ‘pre’ and are the same as the Day 3 values plotted in B. They are re-plotted here for ease of comparison with the post FM1-43 (5 nmol in 2.5 μL) or vehicle (2.5 μL) injection. D) Knee withdrawal threshold of Saline mice before and 90 min after i.a. injection of vehicle (n=2 female; n=2 male). E) Knee withdrawal threshold of Saline mice before and 90 min after i.a. injection of FM1-43 (n=2 female; n=2 male). F) Knee withdrawal threshold of CFA mice before and 90 min after i.a. injection of vehicle (n=2 female; n=2 male). G) Knee withdrawal threshold of CFA mice before and 90 min after i.a. injection of FM1-43 (n=2 female; n=2 male). Paired two-tailed t-tests. B-G) Each dot represents an individual mouse. Female = purple; Male = green.

### FM1-43 inhibits nociceptor responses to mechanical force applied to the knee in an acute inflammatory pain model

To assess whether FM1-43 had direct inhibitory actions on nociceptors *in vivo*, we conducted *in vivo* calcium imaging of the knee-innervating L4 DRG using Na_V_1.8-GCaMP6 mice (Figure 2A). For this experiment we again used the CFA acute knee pain model, as previous studies have shown that local injection of CFA sensitizes sensory neurons as assessed by electrophysiology and calcium imaging [4,7,17,39]. Here, since the mice were expressing GCaMP6s, the red dye FM4-64 was used. During the imaging session, nociceptor responses to 100 g applied to the knee joint were assessed before and 90 min after intra-articular injection of vehicle or FM4-64. CFA mice injected with vehicle had an increase in the number of cells responding post injection to force applied to the knee compared to pre-injection (p=0.0378, Figure 2B-D). Visually, more cells can be seen responding post injection in an imaging area compared to pre at a similar fluorescence intensity (Supplemental Figure 3A-G). In contrast, CFA mice injected with FM4-64 during imaging had a decrease in the number of neurons responding to force applied to the knee compared to pre-injection knee pinch, suggesting that FM4-64 had reduced the responsiveness of nociceptors to the mechanical stimulus (p=0.0016, Figure 2E-G). In a representative imaging area, fewer cells can be seen responding after FM4-64 injection, and those cells still responding appear to respond less intensely (Supplemental Figure 4A-G).

**Figure 2.**
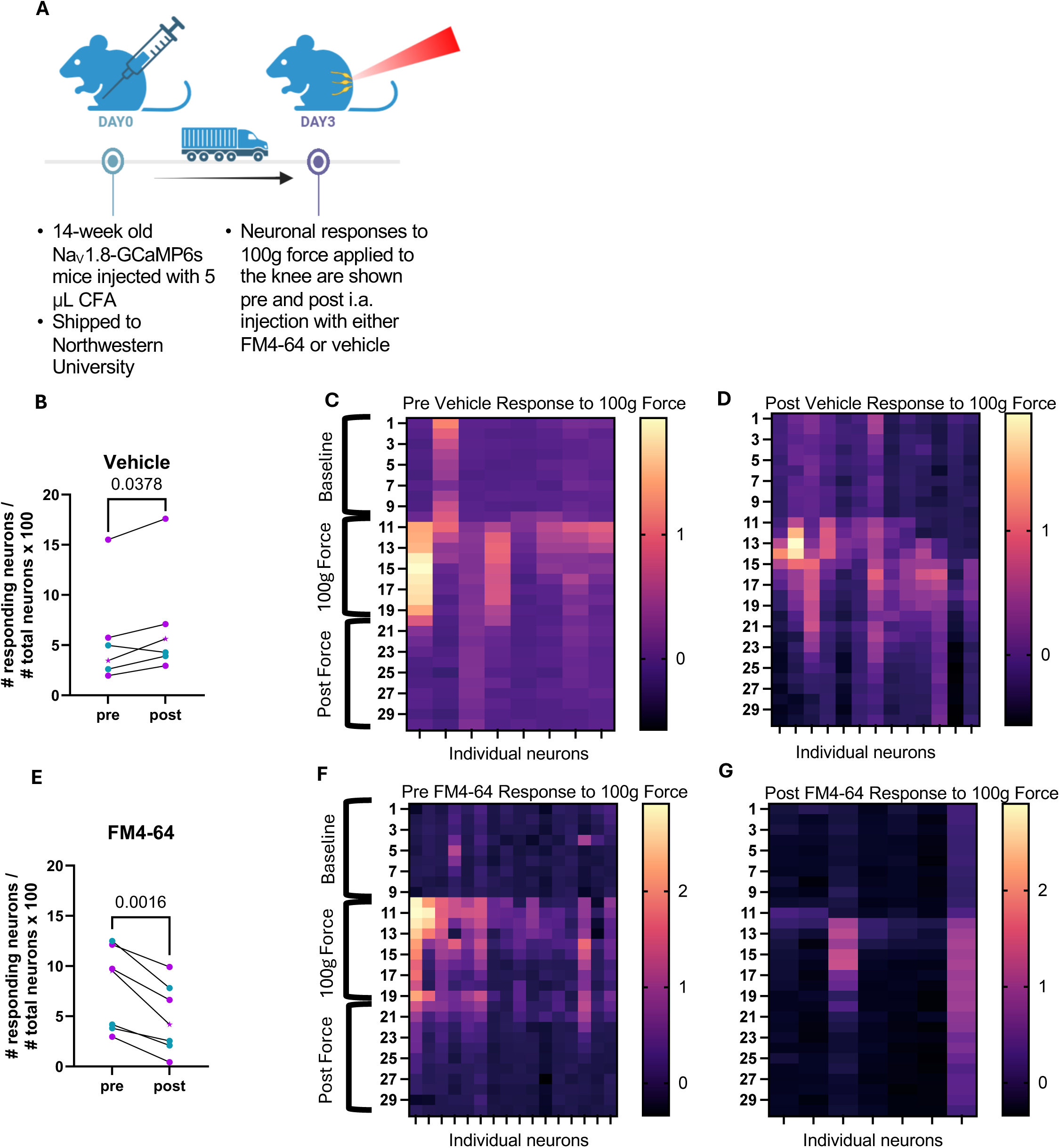
*In vivo* DRG calcium imaging demonstrates that i.a. injection of FM4-64 reduces the number of neuronal responses to force applied to the knee joint of mice 3 days after CFA induction in that knee. A) Experimental design. Mice were injected at Rush University with CFA (5µl stock) then shipped to Northwestern University where *in vivo* calcium imaging occurred 3 days post injection. B) The number of responding neurons in the L4 DRG normalized to the total number of neurons imaged before and after i.a. vehicle injection (5 μL). Paired two-tailed t-test. C,D) Representative heatmaps from the mouse marked with a star in B, depicting the ΔF/Fo values for each frame (row) for each responding neuron (column), Before (C) and 90 min after (D) i.a. vehicle injection. E) The number of responding neurons in the L4 DRG normalized to the total number of neurons imaged before and after i.a. FM4-64 injection (5 nmol in 2.5 μL). Paired two-tailed t-test. F,G) Representative

### FM1-43 demonstrates efficacy in reducing knee hyperalgesia in PMX mice of both sexes

Based on the efficacy shown in the CFA model, we wanted to assess the role of FM1-43 in an OA mouse model, the PMX model, which induces joint damage and pain behaviors in both sexes [15,19,21,29]. To this end we assessed the effectiveness and specificity of FM1-43 in reducing OA pain in male and female WT and Piezo2cko mice subjected to PMX surgery (Figure 3A). By 8 weeks after PMX surgery, WT male and female mice had a lower knee withdrawal threshold compared to sham-operated mice, indicative of knee hyperalgesia (WT PMX: 313 ±9 g; WT Sham: 412±5 g; p<0.0001, Figure 3B). Piezo2cko PMX male and female mice had less knee hyperalgesia compared to WT PMX mice, as we have observed previously in another model of OA [31], but the Piezo2cko mice still had a slightly lower knee withdrawal threshold compared to WT sham mice (Piezo2cko PMX: 390±5 g; p<0.0001 *vs*. WT PMX: 313±9 g; p=0.0664 *vs*. WT sham: 412±5 g, Figure 3B). Intra-articular injection of FM1-43 increased the withdrawal threshold in WT PMX mice compared to pre-injection (pre: 318 ±6 g; post: 372±12 g; p=0.0063, Figure 3C), while saline injection had no effect (pre: 320 ±5 g; post: 319±7 g; p=0.9028, Figure 3D). WT sham mice had less knee hyperalgesia to begin with, but FM injection caused a slight further decrease in knee withdrawal threshold (pre: 419 ±3 g; post: 411±3 g; p=0.1312, Figure 3E), and vehicle injection did as well (pre: 420 ±2 g; post: 411±3 g; p=0.0544, Figure 3F). Unlike WT PMX mice, Piezo2cko PMX mice displayed no change when injected with FM (pre: 393±4 g; post: 396±3 g; p=0.4869, Figure 3G), and injection with saline also had no effect (pre: 389 ±4 g; post: 395 ±3 g; p=0.3274, Figure 3H). These experiments suggest that FM1-43 can reduce local knee mechanical hypersensitivity in a Piezo2-dependent manner.

**Figure 3:**
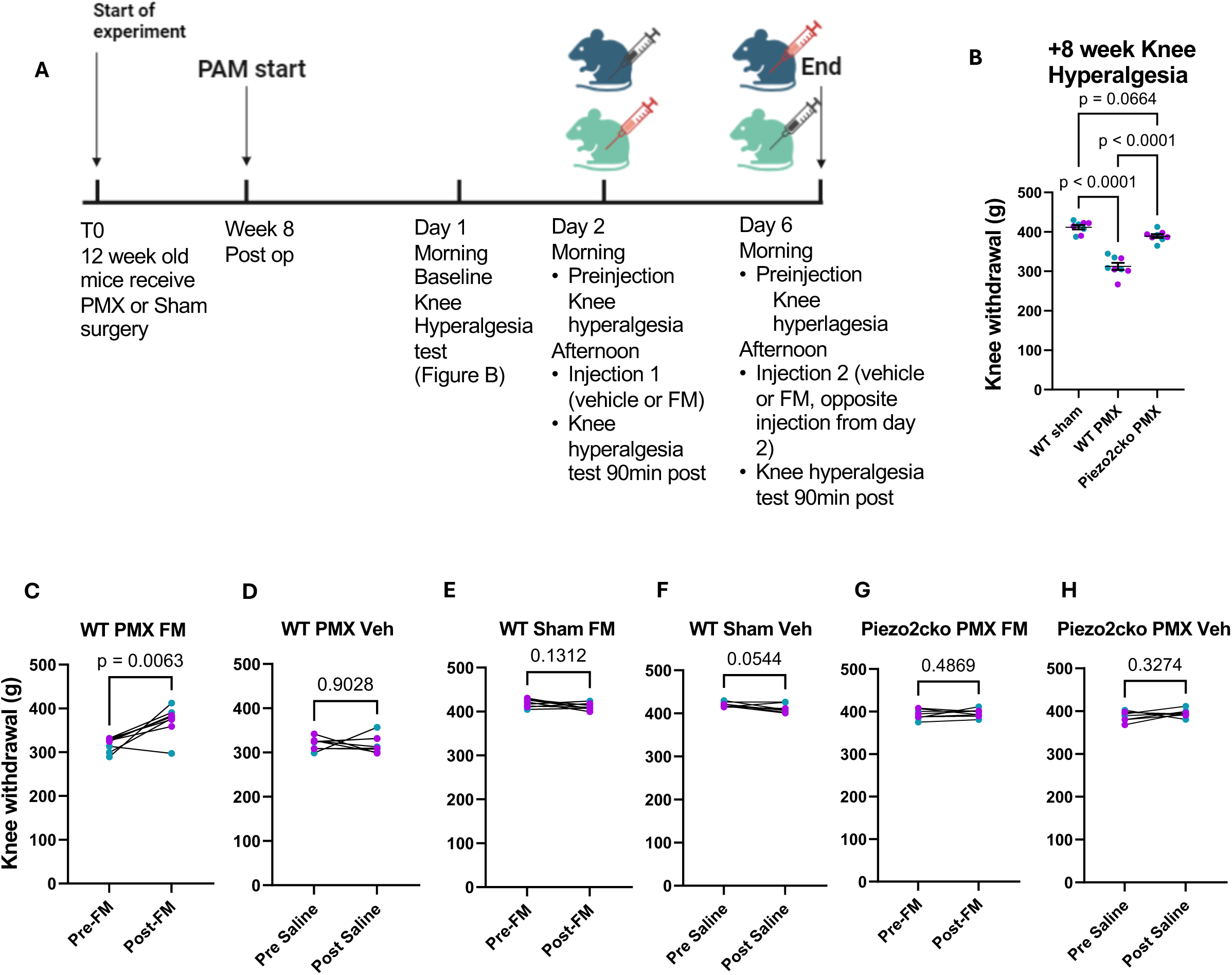
Intra-articular injection of FM1-43 reverses knee hyperalgesia in the PMX model. A) Experimental design. A cross-over study design was used such that mice initially were divided into vehicle and FM1-43 groups for an injection on day 2; on day 6 mice received the opposite injection from day 2. B) Knee withdrawal threshold 8 weeks post PMX or sham surgery. One-way ANOVA with Tukey’s post test. C-H) Plots show data combined from day 2 and day 6 injection tests. C) Knee withdrawal threshold of WT PMX mice before and 90 min after i.a. injection of FM1-43 (n=4 female; n=4 male). D) Knee withdrawal threshold of WT PMX mice before and 90 min after i.a. injection of vehicle (n=4 female; n=4 male). E) Knee withdrawal threshold of WT sham mice before and 90 min after i.a. injection of FM1-43 (n=4 female; n=4 male). F) Knee withdrawal threshold of WT sham mice before and 90 min after i.a. injection of saline vehicle (n=4 female; n=4 male). G) Knee withdrawal threshold of Piezo2cko PMX mice before and 90 min after i.a. injection of FM1-43 (n=4 female; n=4 male). H) Knee withdrawal threshold of Piezo2cko PMX mice before and 90 min after i.a. injection of vehicle (n=4 female; n=4 male). Paired two-tailed t-tests. C-H) Each dot represents an individual mouse. Female = purple; Male = green.

### FM1-43 demonstrates efficacy in reducing weight-bearing asymmetry in PMX mice of both sexes

In a new cohort of PMX mice, we further explored the efficacy of FM1-43 in reducing non-evoked weight-bearing asymmetry, a translationally relevant behavior that develops in the late-stage of the model (>10 weeks after surgery) [21,29]. Twelve weeks after surgery, baseline weight bearing was conducted, and WT male and female mice were randomly assigned to receive vehicle or FM1-43 (Figure 4A). PMX mice placed significantly less weight on their right limb compared to sham mice 12 weeks after surgery (sham: 0.1±0.4 g; PMX: −2±0.3 g; p<0.0001, Figure 4B). When injected with FM1-43, PMX mice placed more weight on their right knee (pre: −3±0.2 g; post: −0.6±0.4 g; p=0.0015, Figure 4C), while vehicle injection did not cause a change in weight bearing (pre: −2±0.3 g; post: −2±0.6 g; p=0.8094, Figure 4D). Sham mice did not display weight-bearing asymmetry to begin with (Figure 4E,F), and FM1-43 injection did not affect weight bearing (pre: 0.1±0.4 g; post: −0.3±0.3 g; p=0.1325 Figure 4E). Likewise, vehicle injection also did not cause a change in weight bearing in sham mice (pre: −0.1±0.4 g; post: −0.1±0.5 g; p>0.9999, Figure 4F).

**Figure 4.**
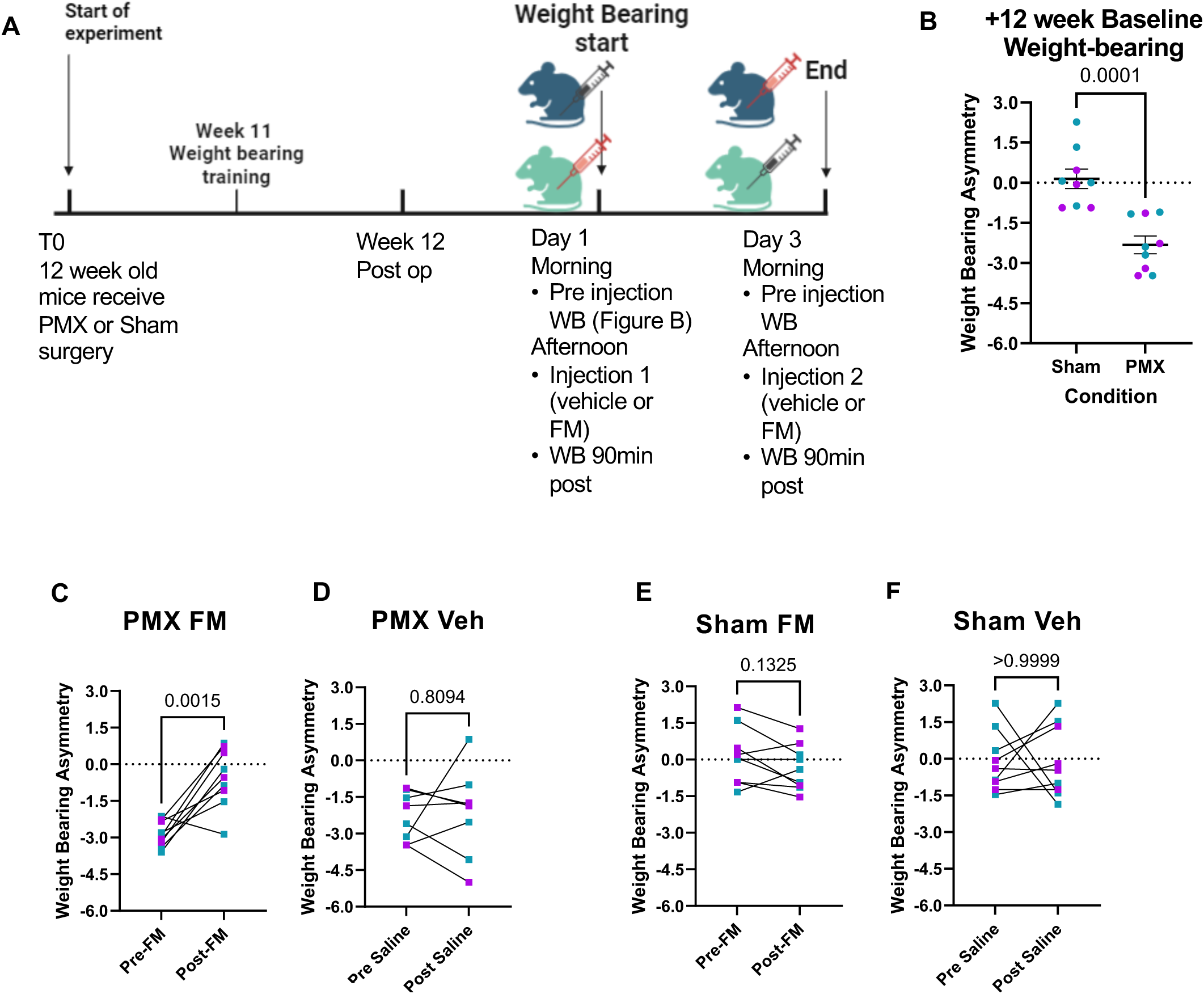
Intra-articular injection of FM1-43 reverses Weight-bearing deficits in the PMX model. A) Experimental timeline. B) Weight bearing asymmetry 12-weeks after WT PMX or sham surgery. Unpaired Student’s t-test. C-F) A cross-over study design was used such that mice initially were divided into vehicle and FM1-43 groups for an injection on day 1; on day 3 mice received the opposite injection from day 1. Plots show data combined from day 1 and day 3 injection tests. C) Weight bearing asymmetry of WT PMX mice before and 90 min after i.a. injection of FM1-43 (n=4 female; n=5 male). D) Weight bearing asymmetry of WT PMX mice before and 90 min after i.a. injection of vehicle (n=4 female; n=5 male). E) Weight bearing asymmetry of WT sham mice before and 90 min after i.a. injection of FM1-43 (n=4 female; n=5 male). F) Weight bearing asymmetry of WT sham mice before and 90 min after i.a. injection of saline vehicle (n=4 female; n=5 male). Paired two-tailed t-tests. C-F) Each dot represents an individual mouse. Female = purple; Male = green.

To assess whether motor coordination of these mice was impacted due to FM1-43 injection, we performed a horizontal ladder test on this set of mice 13 weeks after PMX and sham surgery. There were no significant differences in mice receiving FM1-43 or saline in motor coordination (Supplemental Figure 5), indicating that intra-articular injection of FM1-43 did not cause pronounced proprioceptive deficits.

Histopathology was assessed in the right knees 13 weeks after surgery (Supplemental Figure 6A). As previously demonstrated, male PMX mice had more cartilage damage compared to sham mice (sham: 0±0; PMX: 21±2; p<0.0001, Supplemental Figure 6B) [19,29]. Osteophyte width and synovial pathology were higher in male PMX mice compared to sham as well (sham: 7±7; PMX: 265.2±21; p<0.0001, sham: 0.5±0.3; PMX: 6±0.5; p<0.0001, Supplemental Figure 6C-D). Females overall developed milder pathology compared to males, but PMX surgery still induced increased cartilage damage, osteophyte width, and synovial pathology when compared to female sham mice (sham: 0±0; PMX: 6±1; p=0.0006, sham: 0±0; PMX: 156±10; p<0.0001, sham: 0.5±0.3; PMX: 6±1; p=0.0005, Supplemental Figure 6E-G).

### FM1-43/FM4-64 demonstrates efficacy in reducing knee hyperalgesia associated with spontaneous OA in aged mice of both sexes

Male and female naïve WT and Piezo2cko littermates were aged for 24 months, an age at which they have been shown to develop primary age-associated knee OA[16,31]. At 24 months, WT male mice had significantly more knee hyperalgesia compared to their Piezo2cko littermates (WT: 234±9 g; *vs.* Piezo2cko: 328±8 g; p<0.0001, Figure 5A). After baseline testing, mice were intra-articularly injected with FM1-43 or FM4-64 depending on their genotype and tested 90 minutes later. Mice expressing GCaMP6s were injected with the red FM4-64 dye, and td-Tomato expressing mice were injected with the green FM1-43 dye. FM injection caused a significant increase in knee withdrawal threshold compared to baseline in WT mice, indicating reduced knee hyperalgesia (Pre: 234±9 g; *vs.* Post: 351±27 g; p=0.0112, Figure 5B). FM did not impact the Piezo2cko mice from their baseline score (Pre: 328±8 g; *vs.* Post: 353±8 g; p=0.0866, Figure 5C). Similar to male mice, female WT mice had a significantly lower knee withdrawal threshold when compared to Piezo2cko littermates (WT: 272±21 g; *vs.* Piezo2cko: 343±16 g; p=0.0176, Figure 5D). FM injection also significantly increased the knee withdrawal threshold in female WT mice (Pre: 272±21 g; *vs.* Post: 355±12 g; p=0.0007, Figure 5E), while FM did not significantly impact the knee withdrawal of aged Piezo2cko mice (Pre: 343±16 g; *vs.* Post: 360±7 g; p=0.3050, Figure 5F).

**Figure 5:**
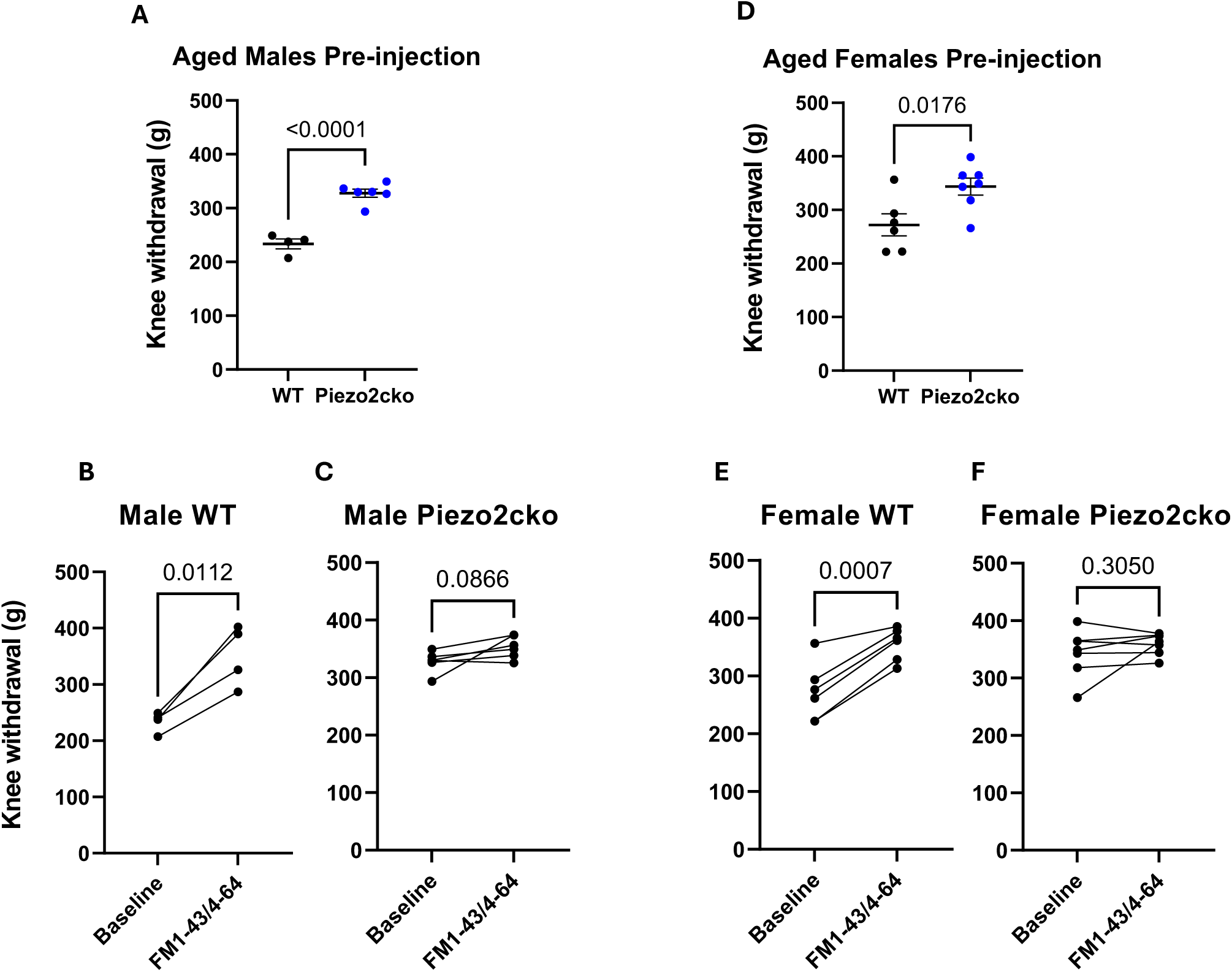
Intra-articular injection of FM dye reverses knee hyperalgesia in aged mice. A) Knee withdrawal threshold of male 24-month-old Piezo2cko and WT mice pre-injection. Student’s two-tailed t-test. B) Knee withdrawal threshold of male WT aged mice pre and post FM. C) Knee withdrawal threshold male Piezo2cko mice pre and post FM. B-C) Paired two-tailed t-test. D) Knee withdrawal threshold of Female 24-month-old Piezo2cko and WT mice pre-injection. Student’s two-tailed t-test. E) Knee withdrawal threshold of female WT aged mice pre and post FM. F) Knee withdrawal threshold female Piezo2cko mice pre and post FM. E-F) Paired two-tailed t-test. Each dot represents an individual mouse.

## Discussion

Targeting Piezo2 via FM1-43 or FM4-64 in acute inflammatory and OA models of knee joint pain reduced knee hyperalgesia and weight-bearing pain behaviors in mice of both sexes. Through *in vivo* calcium imaging of the DRG, we showed that FM-dye injected into the knee joint reduced the responsiveness of nociceptors to mechanical force applied to the knee joint, supporting the proposed mechanism of action that FM-dye acts as a permeant blocker as it is being taken up through active Piezo2 ion channels *in vivo*. Here, we have also confirmed the selectivity of FM-dyes for Piezo2 by demonstrating a lack of effect on knee hyperalgesia in nociceptor-specific Piezo2cko mice. Finally, we demonstrated that local injection of FM1-43 did not cause any observed adverse effects on motor coordination.

Our finding that FM1-43 can reduce acute inflammatory knee pain behavior *in vivo* is consistent with prior literature showing that intraplanar injection of FM1-43 reduced pain behavior caused by an inflammatory agonist [12]. Here, we provide evidence that local administration of FM-dyes to the knee joint can reverse knee hyperalgesia associated with the CFA model of acute knee pain in both male and female mice. Additionally, we demonstrate that intra-articular injection of FM-dyes can transiently reverse knee hyperalgesia and weight-bearing asymmetry associated with models of persistent OA knee pain, the PMX surgical model as well as aging-associated spontaneous OA, in both male and female mice. Together these results support the use of locally administered FM-dyes for inhibition of painful conditions, especially mechanically-driven pain conditions, which may be of relevance for musculoskeletal and rheumatic disease models.

Inflammatory pain caused by local injection of CFA has been well established to induce pain and increase spontaneous neuronal firing as assessed by electrophysiology and calcium imaging [4,7,14,17,31]. FM1-43 is believed to pass through mechanosensitive ion channel pores and inhibit cations such as calcium from entering [8,13,24]. Further, as assessed by *in vitro* electrophysiology, it has been determined that FM1-43 is capable of reducing mechanically activated currents in sensory neurons [8]. Our result contributes to the literature by suggesting FM1-43 inhibits intracellular Ca^2+^ mobilization in nociceptors *in vivo* in response to mechanical stimuli.

Previous literature has suggested a role of Piezo2 in inflammatory pain and sensitization in both humans and in rodents [9,27,32,33]. Our own lab has published findings to this end by demonstrating a reduction in CFA-induced pain behavior in female Piezo2cko mice as well as a reduction in mechanical sensitization and weight-bearing asymmetry in the destabilization of the medial meniscus (DMM) model of OA in male mice [31]. Here, we demonstrated that FM1-43 was effective in reversing knee hyperalgesia in WT male and female mice in the PMX model of OA as well as in spontaneous OA with aging, whereas FM-dye had no effect in Piezo2cko mice, supporting another recent report demonstrating an *in vivo* selectivity of FM1-43 for Piezo2 [36].

Piezo2 has been shown to be expressed by multiple types of sensory neurons, including nociceptors, C-LTMRs, and proprioceptors in rodents and humans [5,9,31,34,38]. Therefore, as has been observed with certain lines of Piezo2 conditional knock-out mice, injection of FM1-43 has the potential to disrupt sensory neuron processes important for proprioception in addition to reducing pain [34,38]. Here, as assessed by the horizontal ladder test, we did not detect any adverse effects on mouse movements following intra-articular injection of FM1-43.

In this proof-of-concept study with FM-dyes, we focused on an intra-articular route of administration, which was shown to have a transient effect on the pain-related behaviors examined, and no immediate structural effects in the joint were noted. Long-term effects of the dye are beyond the scope of the current work, and future work should focus on further development to increase the duration of its effects. However, we have previously demonstrated that Piezo2cko mice develop joint damage comparable to WT mice in both surgical and aging models of OA, providing some evidence that blocking Piezo2 on nociceptors may not be deleterious to the joint, a lingering concern in the field following the anti-NGF clinical trials [20,31].

Targeting Piezo2 pharmacologically has been a critical need in the field of research. Currently, selective options for blocking Piezo2 do not exist. Dyes have a history of being successful in the development of new therapeutics. Prontosil, an azo dye, was used to develop sulfanilamide in the 1940s and its founder was awarded the Nobel prize in Medicine for its discovery [37]. Furthermore, drugs for treating schizophrenia, depression, anxiety, allergies, and malaria were also developed from dyes [37]. Evidence from our study suggests that pharmacological targeting of Piezo2 with FM styryl dyes is effective in reducing both acute inflammatory and persistent OA knee pain suggesting that Piezo2 may represent a target for OA analgesic drugs.

## Acknowledgments

Anne-Marie Malfait has had a consulting role and/or received research funding from Orion and Novartis. The remaining authors declare no competing interests. This work was supported by the National Institutes of Health (NIAMS) grants R01AR077019 (REM), R01AR077019-03S1 (REM), F31AR083277 (NSA), R01AR064251 (AMM, RJM), R01AR060364 (AMM), P30AR079206 (AMM). We would like to acknowledge the Chicago Center on Musculoskeletal Pain (C-COMP) for technical support.

## Figure Legends

**Supplemental Figure 1.** Pilot experiment to establish the dose and timing of FM1-43 i.a. injections. Data demonstrating the dose response and time course of intra-articular FM1-43 reversal of knee hyperalgesia on day 3 of the CFA model in C57BL/6 male mice (n=2 mice per dose). Mean and standard deviation depicted in the graph.

**Supplemental Figure 2.** Baseline comparison across days. A cross-over study design was used such that mice initially were divided into vehicle and FM1-43 groups for an injection on day 2; on day 6 mice received the opposite injection from day 2. A) Knee withdrawal threshold every morning prior to injection WT sham mice. B) Knee withdrawal threshold every morning prior to injection WT PMX mice. C) Knee withdrawal threshold every morning prior to injection Piezo2cko PMX mice. A-C) Repeated measures one-way ANOVA with Tukey’s post test. Each dot represents an individual mouse. Female = purple; Male = green. Star points indicate a mouse receiving FM1-43 that afternoon. Mean and SEM depicted.

**Supplemental Figure 3:** Representative images of a vehicle mouse response pre and post injection. A) Pre-injection baseline. B) Pre-injection response to 100 g mechanical force applied to the knee. C) Line chart of change in fluorescence of responding neurons to force stimulation in B. Frames 0-9: baseline, 10-19: 100g force, 20-30: post stimulation. E) Post-injection baseline. F) Post injection response to 100 g mechanical force applied to the knee. G) Line chart of change in fluorescence of responding neurons to force stimulation in F. Frames 0-9: baseline, 10-19: 100g force, 20-30: post stimulation. Yellow arrows point to examples of responding neurons.

**Supplemental Figure 4:** Representative images of FM4-64 mouse response pre and post injection. A) Pre-injection baseline. B) Pre injection response to 100 g mechanical force applied to the knee. C) Line chart of change in fluorescence of responding neurons to force stimulation in B. Frames 0-9: baseline, 10-19: 100g force, 20-30: post stimulation. E) Post-injection baseline. F) Post injection response to 100 g mechanical force applied to the knee. G) Line chart of change in fluorescence of responding neurons to force stimulation in F. Frames 0-9: baseline, 10-19: 100g force, 20-30: post stimulation. Yellow arrows point to examples of responding neurons.

**Supplemental Figure 5.** Assessment of limb accuracy by horizontal ladder test in WT PMX and Sham mice post vehicle or FM injection. One-way ANOVA with Tukey’s multiple comparisons test. Each dot represents an individual mouse. Female = purple; Male = green. Mean and SEM depicted.

**Supplemental Figure 6.** Knee histology of PMX mice. A) Representative images of H&E and Safranin-O staining of male and female PMX and sham knees. B) Male PMX cartilage degradation score as assessed by Safranin-O staining. C) Male osteophyte width scoring as assessed by H&E stain. D) Male synovial pathology. E) Female PMX cartilage degradation score as assessed by Safranin-O staining. F) Female osteophyte width scoring as assessed by H&E stain. G) Female synovial pathology. B-G) unpaired two-tailed t-test. Each dot represents an individual mouse. Mean and SEM depicted.

