## Supplemental Figures for "FM-dye inhibition of Piezo2 relieves acute inflammatory and osteoarthritis knee pain in mice of both sexes"

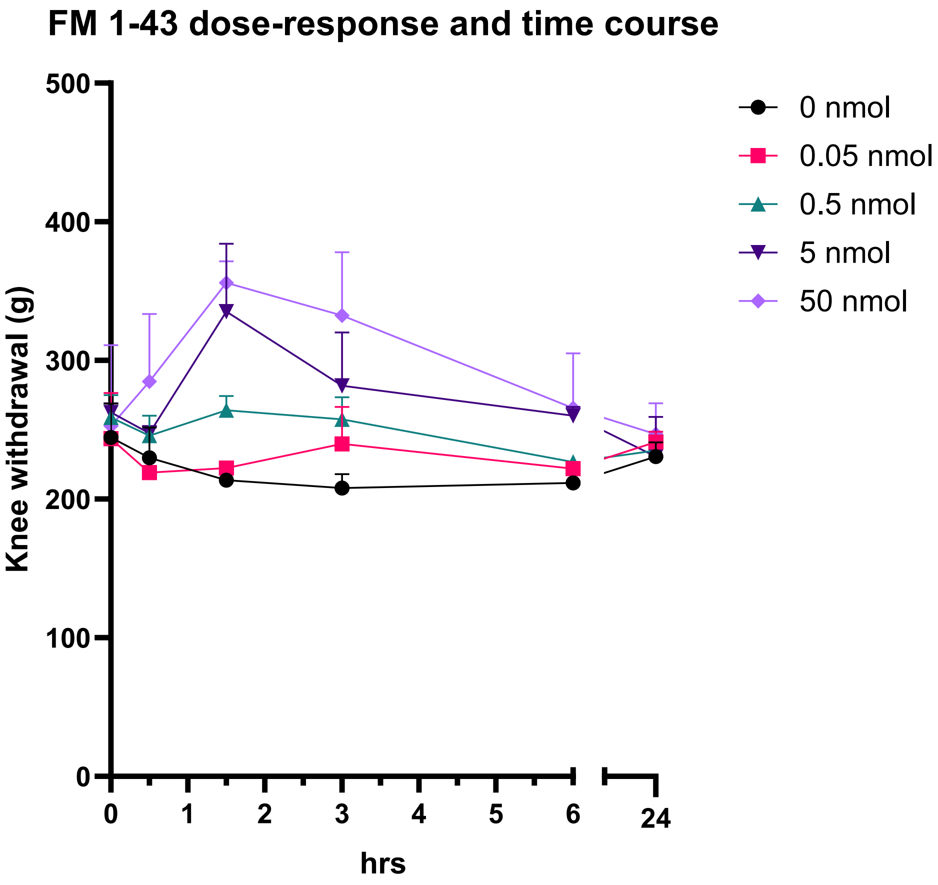


**Supplemental Figure 1.** Pilot experiment to establish the dose and timing of FM1-43 i.a. injections. Data demonstrating the dose response and time course of intra-articular FM1-43 reversal of knee hyperalgesia on day 3 of the CFA model in C57BL/6 male mice (n=2 mice per dose). Mean and standard deviation depicted in the graph.

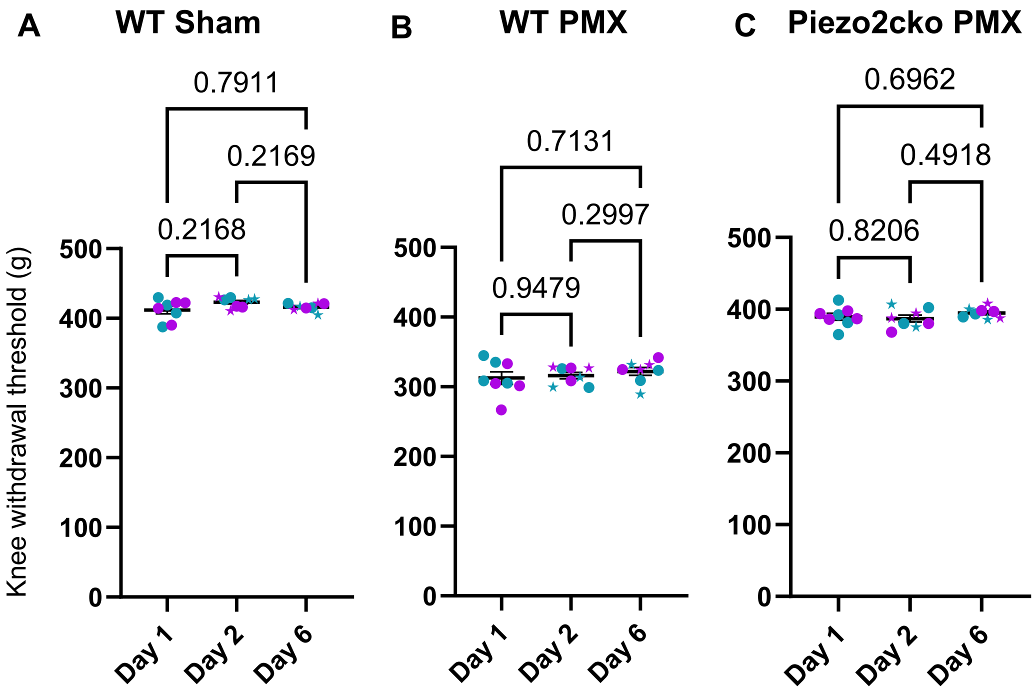


**Supplemental Figure 2**. Baseline comparison across days. A cross-over study design was used such that mice initially were divided into vehicle and FM1-43 groups for an injection on day 2; on day 6 mice received the opposite injection from day 2. A) Knee withdrawal threshold every morning prior to injection WT sham mice. B) Knee withdrawal threshold every morning prior to injection WT PMX mice. C) Knee withdrawal threshold every morning prior to injection Piezo2cko PMX mice. A-C) Repeated measures one-way ANOVA with Tukey's post test. Each dot represents an individual mouse. Female = purple; Male = green. Star points indicate a mouse receiving FM1-43 that afternoon. Mean and SEM depicted.

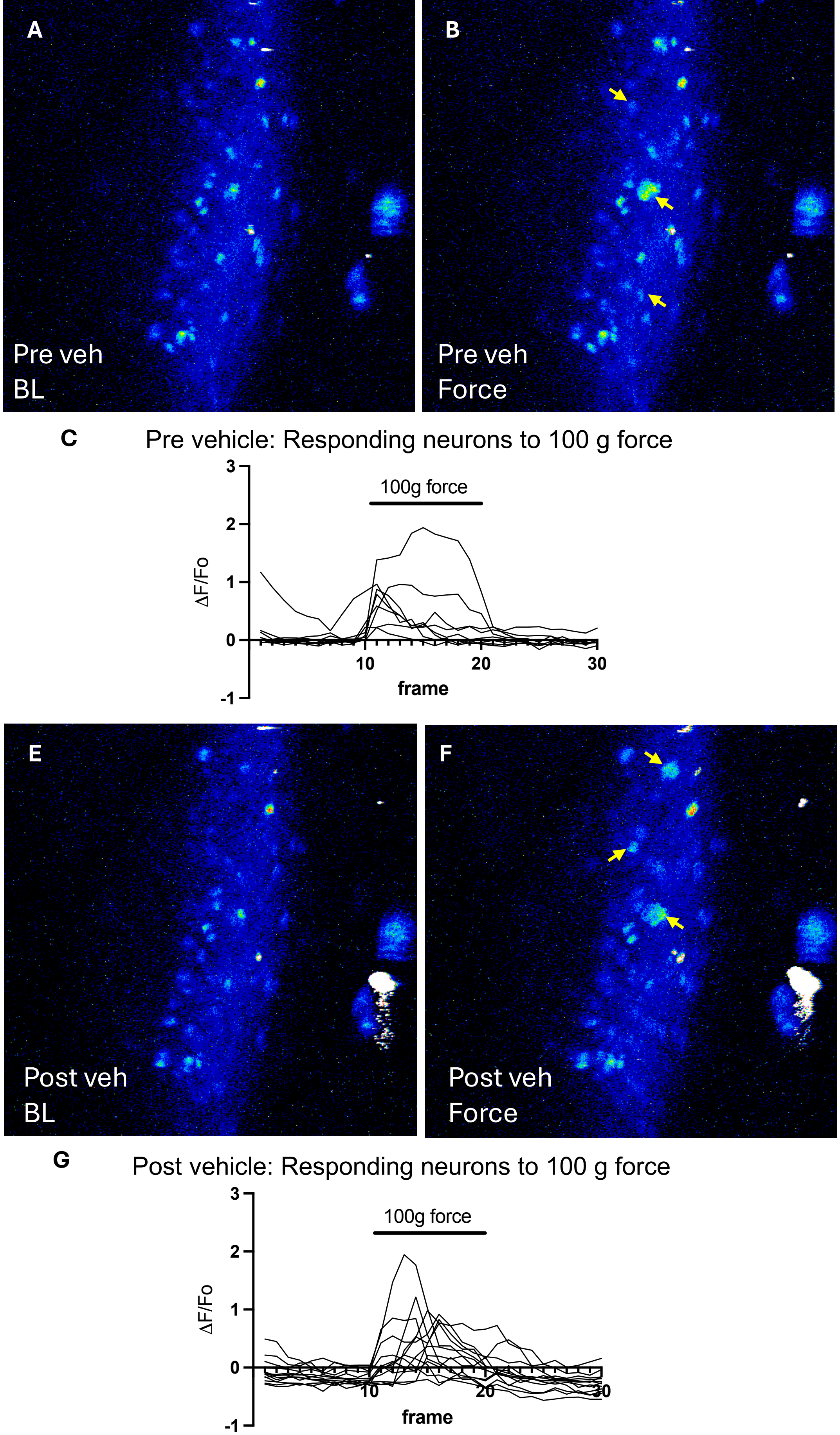


**Supplemental Figure 3**: Representative images of a vehicle mouse response pre and post injection. A) Pre-injection baseline. B) Pre-injection response to 100 g mechanical force applied to the knee. C) Line chart of change in fluorescence of responding neurons to force stimulation in B. Frames 0-9: baseline, 10-19: 100g force, 20-30: post stimulation. E) Post-injection baseline. F) Post injection response to 100 g mechanical force applied to the knee. G) Line chart of change in fluorescence of responding neurons to force stimulation in F. Frames 0-9: baseline, 10-19: 100g force, 20-30: post stimulation. Yellow arrows point to examples of responding neurons.

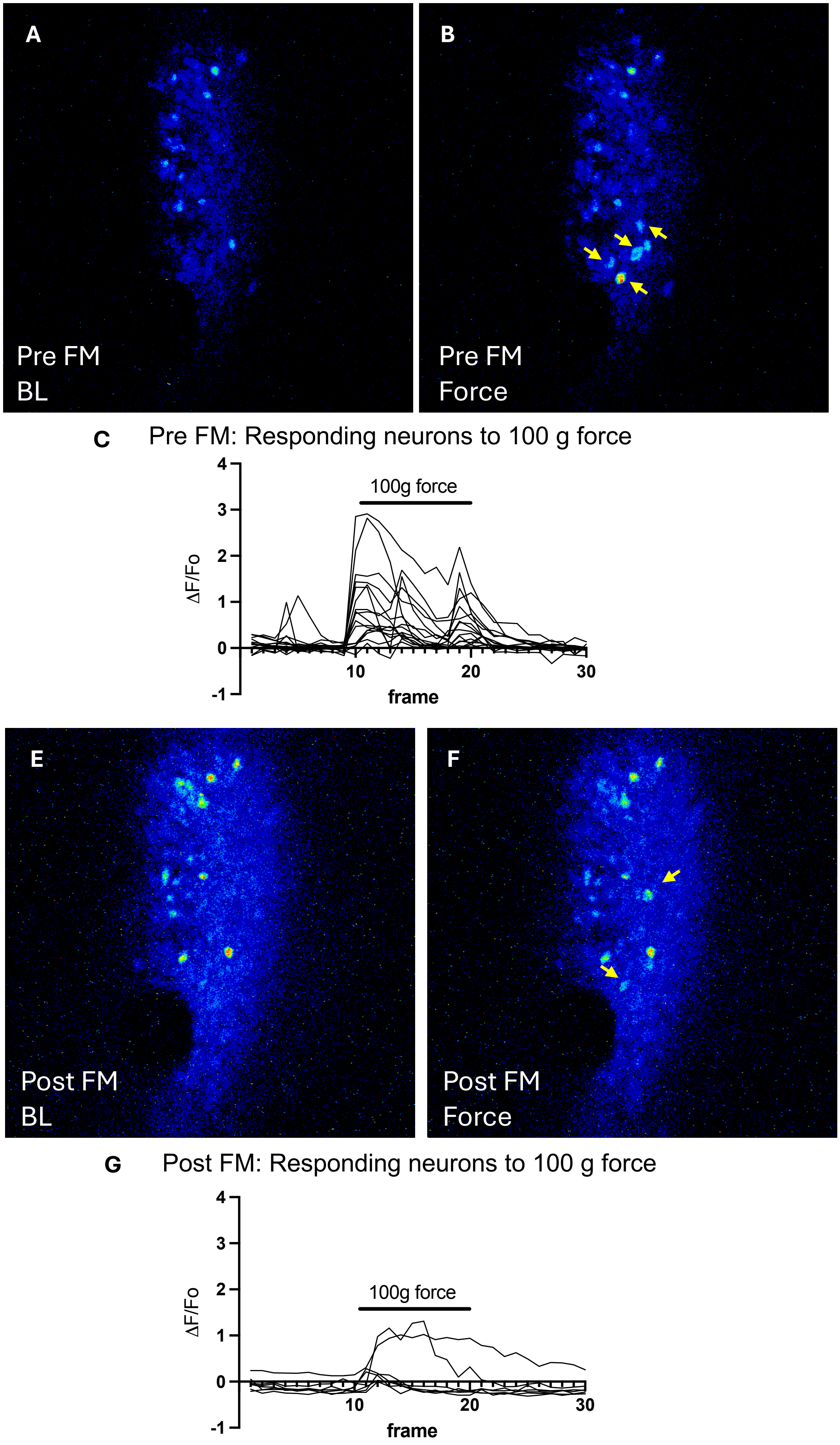


**Supplemental Figure 4**: Representative images of FM4-64 mouse response pre and post injection. A) Pre-injection baseline. B) Pre injection response to 100 g mechanical force applied to the knee. C) Line chart of change in fluorescence of responding neurons to force stimulation in B. Frames 0-9: baseline, 10-19: 100g force, 20-30: post stimulation. E) Post-injection baseline. F) Post injection response to 100 g mechanical force applied to the knee. G) Line chart of change in fluorescence of responding neurons to force stimulation in F. Frames 0-9: baseline, 10-19: 100g force, 20-30: post stimulation. Yellow arrows point to examples of responding neurons.

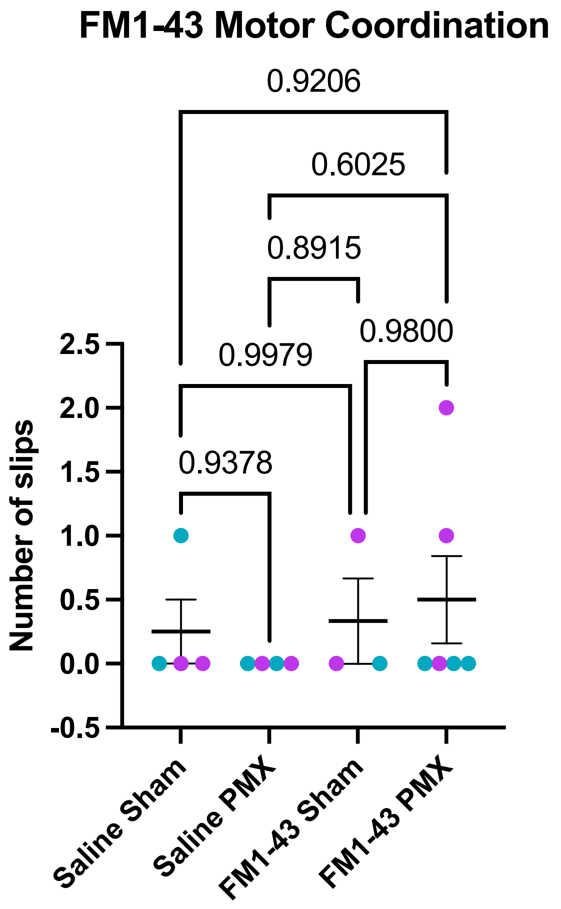


**Supplemental Figure 5**. Assessment of limb accuracy by horizontal ladder test in WT PMX and Sham mice post vehicle or FM injection. One-way ANOVA with Tukey's multiple comparisons test. Each dot represents an individual mouse. Female = purple; Male = green. Mean and SEM depicted.

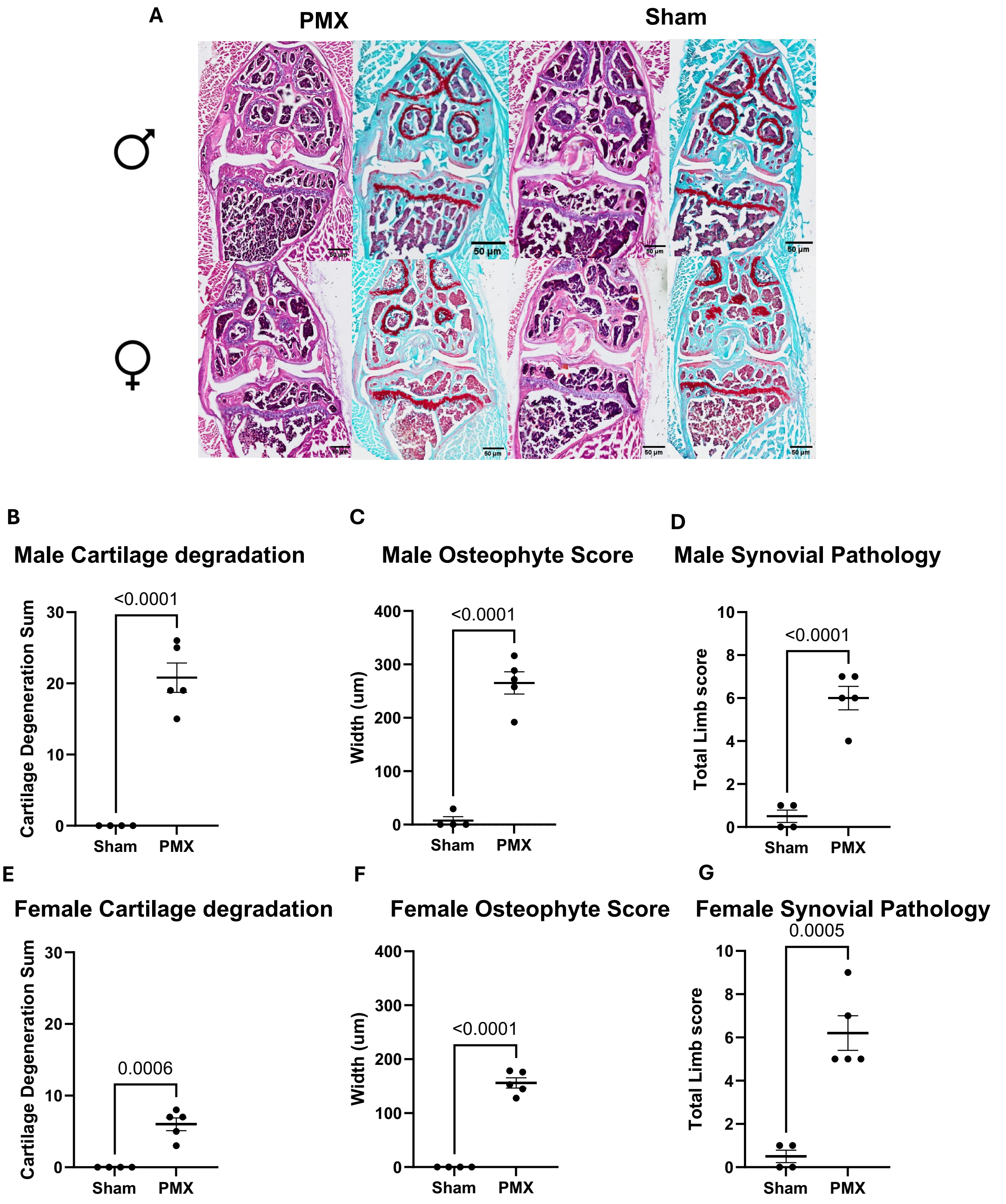


**Supplemental Figure 6**. Knee histology of PMX mice. A) Representative images of H&E and Safranin-O staining of male and female PMX and sham knees. B) Male PMX cartilage degradation score as assessed by Safranin-O staining. C) Male osteophyte width scoring as assessed by H&E stain. D) Male synovial pathology. E) Female PMX cartilage degradation score as assessed by Safranin-O staining. F) Female osteophyte width scoring as assessed by H&E stain. G) Female synovial pathology. B-G) unpaired two-tailed t-test. Each dot represents an individual mouse. Mean and SEM depicted.
